# Dissociable Stimulus- and Response-Driven Serial Dependence Effects in Auditory, Visual, and Audiovisual Motion Judgments

**DOI:** 10.64898/2026.09.17.752446

**Authors:** Adam J. Tiesman, Kalina Stoyanova, Hannah Bertisch, Ramnarayan Ramachandran, Mark T. Wallace

## Abstract

Efficient motion perception requires integrating immediate sensory evidence with internal representations shaped by experience. This influence of recent history can produce serial dependence effects (SDEs), in which current judgments are biased toward or away from the source of previous information. We examined SDEs in auditory, visual, and cued audiovisual motion-direction discrimination using aggregate psychometric measures and trial-level generalized linear mixed-effects models (GLMMs), which estimated the independent contributions of the previous stimulus direction and the participant’s previous response. Participants completed unisensory (n=57) and cued audiovisual (n=48) motion-discrimination tasks. Aggregate analyses indicated attractive SDEs in auditory conditions, repulsive SDEs in visual conditions, and no net SDE in the audiovisual-cued condition. Trial-level modeling revealed a more nuanced structure: previous stimulus direction produced repulsive effects in both unisensory conditions, with previous visual direction influencing judgments across all conditions containing visual motion, whereas previous response produced attractive effects only when visual information was task-relevant. In the audiovisual-cued condition, the absence of a net psychometric shift was consistent with the opposing influences of an attractive response-history effect and a repulsive visual stimulus-history effect rather than an absence of serial dependence. These findings suggest that serial dependence in motion perception reflects dissociable and concurrently operating processes, including a predominantly visual, adaptation-like stimulus-history effect and a task-dependent, decision-related response-history effect.

## 1 Introduction

Efficient interaction with the environment requires combining current sensory evidence with internal representations formed through prior experience ([Shams and Beierholm, 2022, Rohe and Noppeney, 2015, Odegaard et al., 2015]). Within a Bayesian framework, sensory evidence is weighted according to its reliability and combined with prior information to generate a stable perceptual estimate ([Shams and Beierholm, 2022, Rohe and Noppeney, 2015, Ernst and Banks, 2002, Kayser and Heuer, 2024]). Because natural environments typically provide information through multiple senses, this process also involves integrating redundant cues across different sensory modalities. When auditory and visual signals describe the same event, the nervous system combines them according to their relative reliability, producing a more precise estimate than either modality alone ([Ernst and Banks, 2002, Alais and Burr, 2004, Battaglia et al., 2003, Morgan et al., 2008, Murray and Wallace, 2011]). Perception therefore reflects not only current multisensory evidence, but also prior experience, and task demands.

One form of prior information is the perceptual and decisional history established across successive trials. Serial dependence effects (SDEs) are systematic influences of recent stimuli, percepts, or responses on current behavior ([Pascucci et al., 2023, Cicchini et al., 2024]). These effects are often attractive, such that judgments are biased toward the source of preceding information, potentially promoting perceptual stability by integrating related inputs over time ([Fischer and Whitney, 2014, Cicchini et al., 2014, Van der Burg et al., 2021, Kiyonaga et al., 2017]). Under other conditions, recent experience can also produce repulsive effects, in which perception is biased away from the source of previous stimulation ([Fritsche et al., 2017, Alais et al., 2017]). Attractive and repulsive SDEs may therefore reflect dissociable processes: sensory adaptation may produce repulsion away from the previous stimulus, whereas persistence of a previous perceptual estimate, decision criterion, or response may produce attraction toward a preceding choice ([Kohn, 2007, Glasser et al., 2011, Fritsche et al., 2017, Togoli et al., 2026]). Because previous stimuli and responses are typically correlated, these processes likely operate concurrently and can be difficult to distinguish when trial-history factors are examined separately. Notably, recent large-scale evidence indicates that serial dependence does not invariably improve perceptual decisions and can instead increase perceptual error, suggesting that the functional consequences of history effects may depend on which underlying process dominates in a given context ([Ozkirli et al., 2026]).

Motion perception provides an informative context in which to test these competing influences. Because motion depends on change over time, temporal averaging could reduce sensitivity to changes in direction or velocity, making repulsive adaptation particularly useful ([Kohn, 2007, Huk et al., 2001]). Consistent with this account, motion aftereffects occur when exposure to motion in one direction biases subsequent perception in the opposite direction, including following relatively brief exposure ([Glasser et al., 2011, Neelon and Jenison, 2004, Huk et al., 2001]). Recent evidence suggests that serial dependence may arise through partially dissociable, modality-specific mechanisms, with auditory and visual serial dependence exhibiting distinct patterns of feature and spatial selectivity ([Togoli et al., 2026, Hashimoto and Makioka, 2026, Fornaciai et al., 2026]). Visual motion generally provides greater spatial precision than auditory motion, with auditory motion processing relying more on duration and displacement and visual motion processing relying more on velocity ([Freeman et al., 2014, Schoenhaut et al., 2025, Tiesman et al., 2026]). These differences are consistent with the Modality Appropriateness Hypothesis, according to which vision is typically more reliable for spatial judgments and audition is more reliable for temporal judgments ([Welch and Warren, 1980]). Recent work further demonstrates that auditory motion can systematically bias visual motion perception, reinforcing the view that audiovisual motion processing reflects interactions between modality-specific motion representations rather than independent processing streams ([Harada et al., 2026]).

These modality differences generate distinct but compatible predictions. An adaptation-based account predicts a repulsive influence of previous stimulus direction, particularly for the more precise and spatially tuned visual motion representation ([Kohn, 2007, Glasser et al., 2011]). In contrast, Bayesian reliability weighting predicts greater reliance on prior information when sensory evidence is uncertain, potentially producing stronger attractive history effects in audition ([Ernst and Banks, 2002, Rohe and Noppeney, 2015]). Such attraction could arise from previous sensory estimates or from the persistence of previous decisions and responses, and its expression may depend on which modality is currently guiding behavior ([Fritsche et al., 2017]). Serial dependence is therefore likely to reflect concurrent stimulus- and response-history processes whose relative contributions vary across modality and attentional context.

The present study examined SDEs in auditory (A), visual (V), and cued audiovisual (AV) motion-direction discrimination (see Figure 1 for trial structure). Psychometric functions were used to characterize history-dependent changes in point of subjective equality (PSE), providing a measure of net history effects. Trial-level generalized linear mixed-effects models were then used to estimate the independent contributions of previous stimulus direction and previous response while accounting for current sensory evidence and other trial-level factors. We predicted that previous stimulus direction would produce repulsive effects, particularly for visual motion, while response-related processes could produce attractive effects. More broadly, we hypothesized that motion serial dependence would not be captured by a single auditory-versus-visual or attractive-versus-repulsive distinction, but would instead reflect separable stimulus- and response-history influences that may reinforce or oppose one another across conditions.

**Figure 1.**
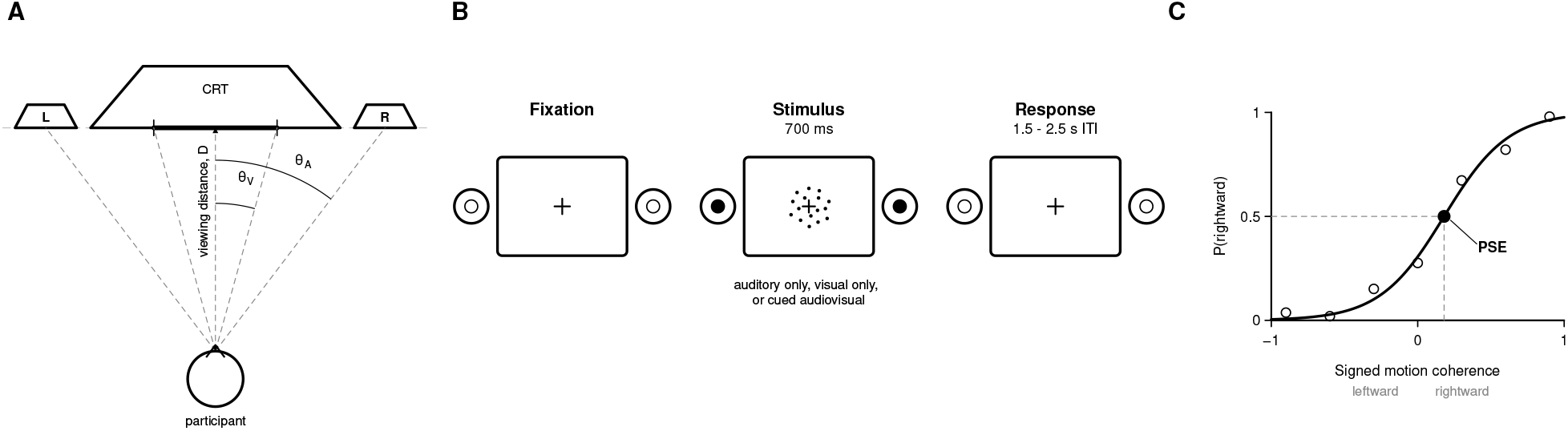
Experimental Overview. **(A)** Participants were seated in front of a CRT monitor flanked with two speakers. **(B)** For all conditions, participants were instructed to fixate on a fixation cross in the middle of the screen, followed by a 700 ms A, V, or AV stimulus presentation and ending with a jittered response window. **(C)** Behavioral responses were fit to psychometric curves, and the point of subjective equality (PSE) was used as a measure of leftward/rightward response bias.

## 2 Results

### 2.1 SDEs in PSEs Based on Prior Response

To investigate whether the responses in the prior trial influenced response bias (PSE) in the current trial, psychometric curves were fit separately to the set of trials following a leftward response and the set of trials following a rightward response, and the resulting PSEs were compared with each other. Critically, an SDE is defined not by whether either PSE alone differs from zero, but by whether the PSE following leftward responses differs from the PSE following rightward responses: a difference in which responses are biased toward the direction of the previous response constitutes an attractive SDE, while a difference in which responses are biased away from the previous response direction constitutes a repulsive SDE (Figure 1C). At the group level, this comparison was made by calculating ΔPSE obtained by subtracting the PSE after right responses from the PSE after left responses for each subject. The resultant ΔPSE was then tested against zero using a Wilcoxon signed-rank test. Positive ΔPSE indicate an attractive SDE while negative values indicate a repulsive SDE (Figure 2).

**Figure 2.**
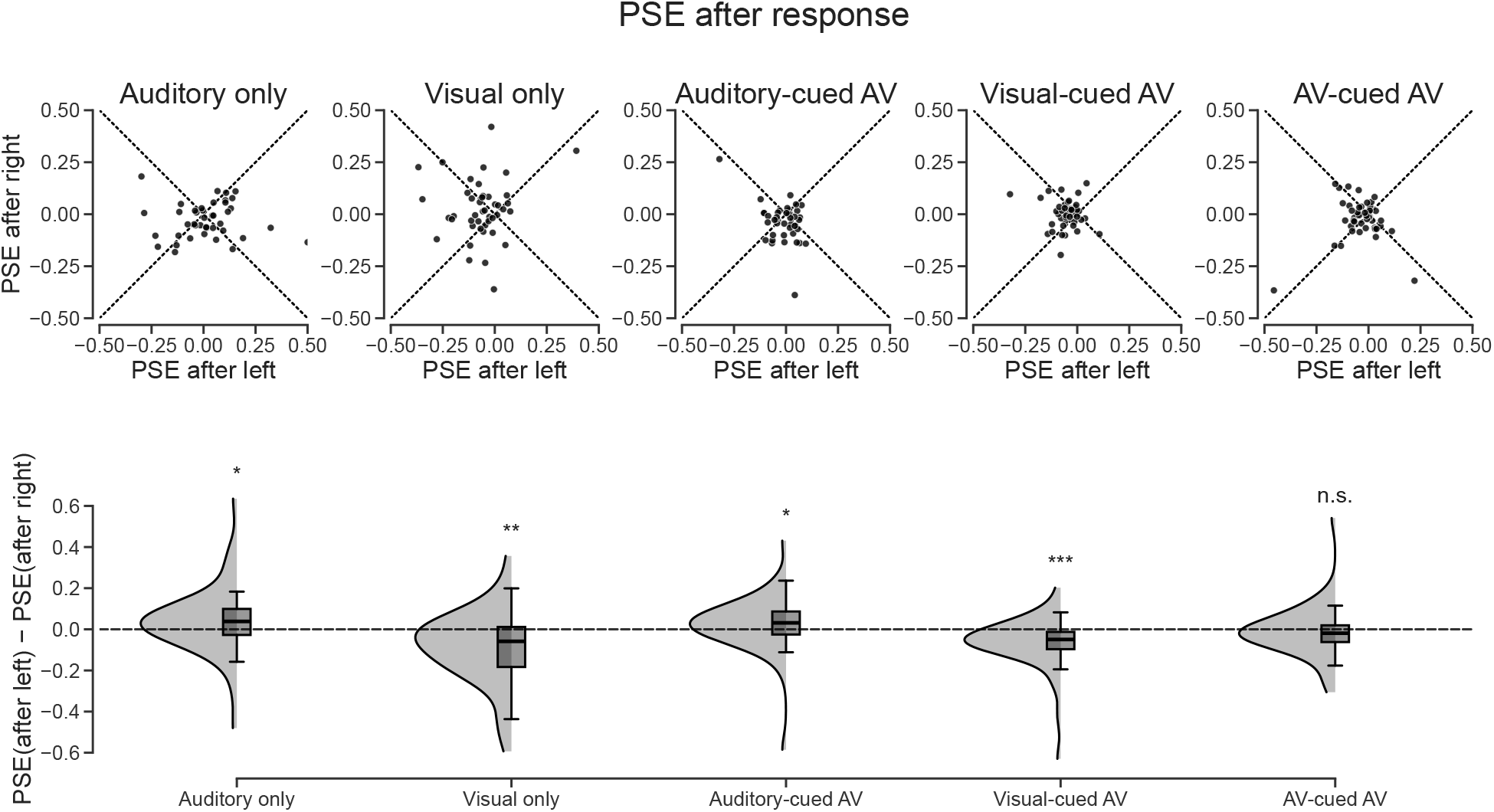
Serial dependence in response bias based on the prior trial’s response. **Top row:** Point of subjective equality (PSE) following rightward responses plotted against PSE following leftward responses, shown separately for each of the five task conditions. Each dot represents an individual participant. Points below the diagonal line of equality indicate a repulsive SDE (bias away from the previous response direction); points above indicate an attractive SDE (bias toward the previous response direction). **Bottom row:** Distribution of the resulting ΔPSE (PSE after left *−*PSE after right) for each condition, shown as kernel density estimates with overlaid box plots. Positive values indicate an attractive SDE and negative values a repulsive SDE; the dashed line marks zero (no SDE). Significance markers refer to Wilcoxon signed-rank tests of ΔPSE against zero: * *p <* 0.05, ** *p <* 0.01, *** *p <* 0.001, n.s. *p >* 0.05.

In the auditory-only condition, the mean ΔPSE was significantly greater than zero (Wilcoxon signed-rank test: *p* = 0.050), indicating that responses exhibited an attractive SDE (Figure 2). In contrast, in the visual-only condition, the ΔPSE was significantly less than zero (Wilcoxon signed-rank *p* = 0.002), indicating a repulsive SDE (Figure 2). Thus, in the unisensory conditions, auditory-only motion judgments were, on the average, attractive and visual-only motion judgments were, on the average, repulsive to the responses in the previous trial.

A similar pattern held for the cued audiovisual conditions. In the auditory-cued AV condition, the mean ΔPSE was significantly greater than zero (Wilcoxon signed-rank test: *p* = 0.045), indicating that responses exhibited an attractive SDE (Figure 2). In contrast, visual-cued AV ΔPSE was significantly less than zero (Wilcoxon signed-rank *p <* 0.001), indicating a repulsive SDE (Figure 2). Interestingly, ΔPSE did not significantly differ from zero in the audiovisual-cued AV condition (*p* = 0.219; Figure 2), indicating no net PSE-based SDE. This result could reflect the summation (and consequent cancellation) of the attractive and repulsive SDEs seen in the auditory-cued and visual-cued conditions, respectively.

### 2.2 SDEs in PSEs Based on Prior Direction

A similar approach was used to test whether the stimulus direction in the prior trial, rather than the participant’s response, influenced PSE. In the auditory-only condition, the mean ΔPSE (for trials after a rightward relative to a trial after a leftward stimulus) was significantly greater than zero (Wilcoxon signed-rank test: *p* = 0.003), indicating that responses exhibited an attractive SDE for stimulus-direction history (Figure 3). In contrast, visual-only ΔPSE did not significantly differ from zero (Wilcoxon signed-rank *p* = 0.351), indicating no SDE based on prior stimulus direction in vision (Figure 3).

**Figure 3.**
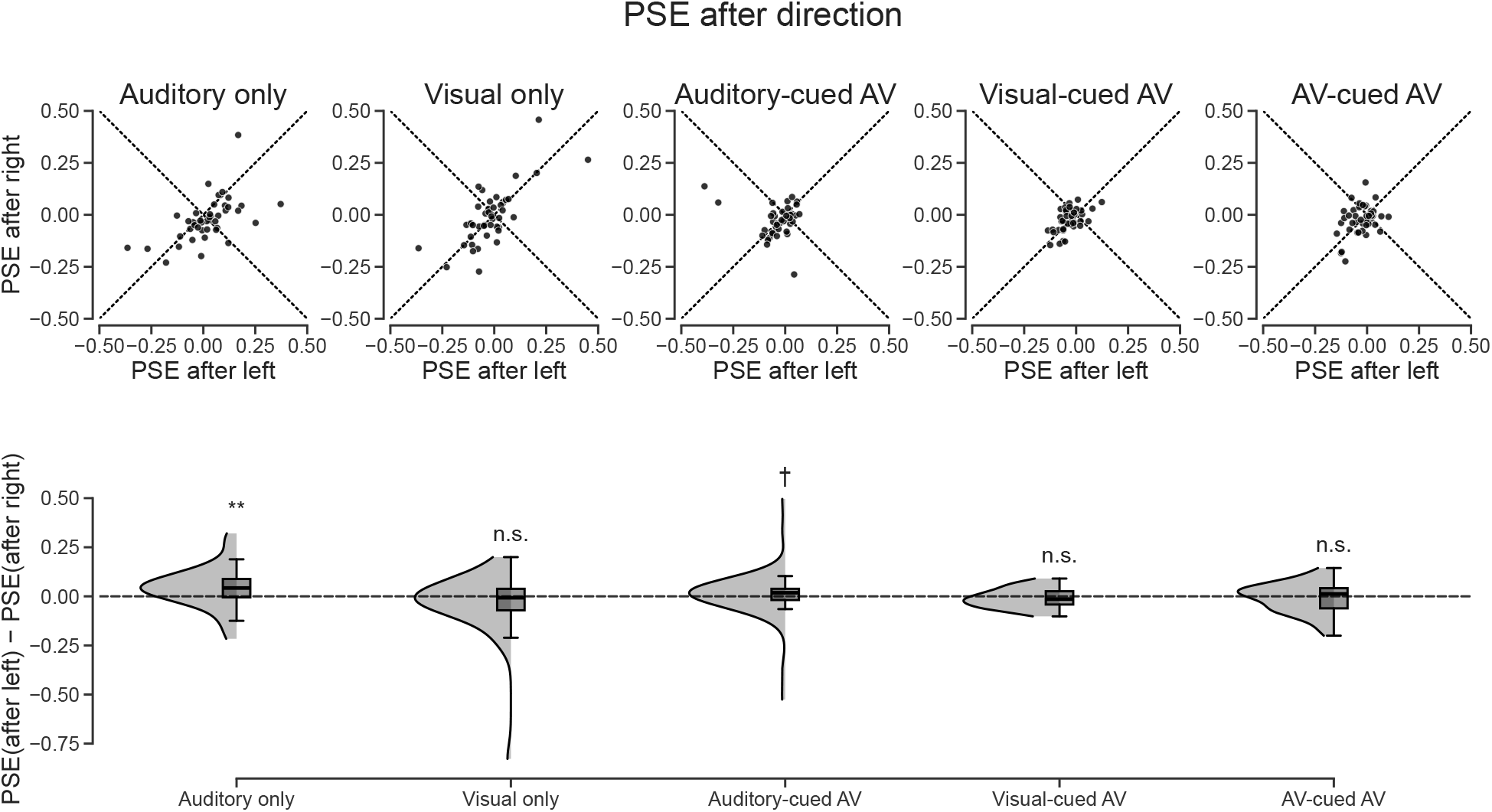
Serial dependence in response bias based on the prior trial’s stimulus direction. Conventions are identical to Figure 2. **Top row:** PSE following rightward stimulus directions plotted against PSE following leftward stimulus directions, for each of the five task conditions, with each dot representing an individual participant. **Bottom row:** Distribution of ΔPSE (PSE after left *−*PSE after right) per condition, with positive values indicating an attractive SDE and negative values a repulsive SDE. Significance markers refer to Wilcoxon signed-rank tests of ΔPSE against zero: *†p <* 0.10, * *p <* 0.05, ** *p <* 0.01, *** *p <* 0.001, n.s. *p >* 0.10.

The same analysis was performed for the cued audiovisual conditions. In the auditory-cued condition, the mean ΔPSE was trending greater than zero (Wilcoxon signed-rank test: *p* = 0.067), indicating that responses tended towards an attractive SDE for stimulus-direction history (Figure 3). In contrast, ΔPSE did not significantly differ from zero in either the visual-cued (*p* = 0.192) or audiovisual-cued (*p* = 0.663) conditions, indicating no stimulus-direction-based SDE in either (Figure 3). Overall, this indicates that stimulus attributes (i.e., the sensory information) produced weaker SDEs compared to the previous response.

### 2.3 Trial-Level Generalized Linear Mixed-Effects Modeling of Motion Judgments

These SDE results reflected a potential contribution of both current sensory evidence and trial history to a given response. Because PSE measures reflect responses aggregated over a given condition, we sought to interrogate these effects at the single trial level. To do this, we aggregated all trials across participants (Experiment1 : *N* = 57 participants, *N* = 19,404 trials; Experiment 2: *N* = 33 participants, *N* = 22,736 trials) and built a generalized linear mixed model (GLMM) for the probability of a rightward response (see Methods, section 4.5). Rather than fitting separate models within each participant, we fit a single model pooling trials across participants with participant-level random effects. Because we hypothesized history effects would be modest relative to current sensory evidence, per-participant fits should be poorly constrained by the trials available from any one individual, whereas pooling estimates these coefficients from the full trial set while random effects accommodate individual differences. We first fit a reduced model containing only current sensory evidence, together with nuisance covariates for unsigned coherence and trial number, to confirm that evidence reliably predicted trial-level choice before evaluating trial history. This model was fit separately for the unisensory experiment (auditory-only, visual-only; auditory-only as reference) and the cued audiovisual experiment (auditory-cued AV, visual-cued AV, audiovisual-cued AV; auditory-cued AV as reference).

Current sensory evidence significantly predicted the probability of a rightward response in every condition of both experiments (all *ps <* .001), with one exception: auditory evidence did not significantly predict choice during visual-cued AV trials (*β* = 0.13, *p* = .783), consistent with an unattended modality carrying little weight when a different modality is explicitly cued (Table 2). In the unisensory experiment, this slope was steeper for visual than auditory evidence (Visual *−*Auditory contrast: *β* = 0.76, *p <* .001; Table 3), consistent with the higher visual discrimination sensitivities reported ([Tiesman et al., 2026]). In the cued AV studies, visual evidence remained numerically the stronger predictor of choice even during auditory-cued AV trials, again paralleling the sensitivity patterns reported ([Tiesman et al., 2026]).

We next tested whether adding trial-history predictors — the participant’s previous response, the direction of the previous trial’s stimulus, the previous trial’s coherence, and whether the previous trial was a catch trial — improved model fit relative to the evidence-only model. For both experiments, the full history model fit significantly better than the reduced model (unisensory: *χ*^2^(6) = 143.51, *p <* .001, ΔAIC = 131.5, ΔBIC = 84.3; cued audiovisual: *χ*^2^(11) = 174.29, *p <* .001, ΔAIC = 152.3, ΔBIC = 63.9), indicating that trial history captured reliable variance in choice beyond current sensory evidence alone.

Trial-history effects were modality- and condition-specific (Table 1, Table 2). In the unisensory experiment, the participant’s previous response was a significant positive (attractive) predictor of the current response in the visual-only condition (*β* = 0.69, *p <* .001) but not in the auditory-only condition (*β* = *−*0.08, *p* = .564); this difference between modalities was itself significant (*β* = 0.77, *p <* .001; Table 3). In contrast, the direction of the previous trial’s stimulus was a significant negative (repulsive) predictor of the current response in both the auditory-only (*β* = *−*0.23, *p <* .001) and visual-only conditions (*β* = *−*0.21, *p* = .001), with no significant difference between them (*β* = 0.02, *p* = .761; Table 3).

**Table 1.** Unisensory GLMM: current-evidence weight and trial-history effects by condition. Estimates are GLMM log-odds coefficients (95% CI), testing whether each is significantly different from zero. N=57 participants, 19404 trials. *p*-values are uncorrected (pre-specified, condition-specific tests; history terms are gated by the significant history-vs-no-history likelihood-ratio test reported in the main text).

| Condition | Term | Estimate [95% CI] | $p$ | Sig. |
| --- | --- | --- | --- | --- |
| Auditory only | Current evidence | 5.241 [4.144, 6.339] | $< 0.001$ | *** |
| Visual only | Current evidence | 6.002 [4.900, 7.104] | $< 0.001$ | *** |
| Auditory only | Previous response | -0.078 [-0.344, 0.187] | 0.564 | n.s. |
| Visual only | Previous response | 0.694 [0.427, 0.961] | $< 0.001$ | *** |
| Auditory only | Previous stimulus direction | -0.230 [-0.351, -0.109] | $< 0.001$ | *** |
| Visual only | Previous stimulus direction | -0.206 [-0.331, -0.082] | 0.001 | ** |

In the cued audiovisual experiment, previous-response effects again depended on cue condition, but in a more complex pattern: previous response was a significant **positive** predictor of choice during visual-cued AV (*β* = 0.66, *p <* .001) and audiovisual-cued AV trials (*β* = 0.27, *p <* .001), but a significant **negative** predictor during auditory-cued AV trials (*β* = *−*0.34, *p <* .001) — all three conditions showed a reliable effect, but auditory-cued AV trials showed the opposite sign from the other two (Table 2). The direction of the previous trial’s visual stimulus was a consistent negative predictor of choice across all three cued audiovisual conditions (*β* = *−*0.19 to *−*0.40, all *ps ≤* .015; Table 2), whereas the direction of the previous trial’s auditory stimulus did not reliably predict choice in any cue condition (all *ps >* .16). Notably, in the audiovisual-cued AV condition — the one condition that showed no net PSE shift in the classical analysis above (section 2.1, Figure 2) — the GLMM identified two significant, opposing history effects (a positive previous-response effect and a negative previous-visual-direction effect) rather than a true absence of history-dependent bias, suggesting the null PSE shift in this condition may reflect the summation of opposing effects rather than an absence of serial dependence.

**Table 2.** Cued-AV GLMM: current-evidence weight and trial-history effects by cue condition. Estimates are GLMM log-odds coefficients (95% CI), testing whether each is significantly different from zero. N=33 participants, 22736 trials. *p*-values are uncorrected (pre-specified, condition-specific tests; history terms are gated by the significant history-vs-no-history likelihood-ratio test reported in the main text).

| Condition | Term | Estimate [95% CI] | $p$ | Sig. |
| --- | --- | --- | --- | --- |
| Auditory-cued AV | Auditory evidence | 3.919 [3.105, 4.734] | < 0.001 | *** |
| Visual-cued AV | Auditory evidence | 0.127 [-0.781, 1.036] | 0.783 | n.s. |
| AV-cued AV | Auditory evidence | 2.358 [1.493, 3.224] | < 0.001 | *** |
| Auditory-cued AV | Visual evidence | 4.008 [3.014, 5.002] | < 0.001 | *** |
| Visual-cued AV | Visual evidence | 10.827 [9.751, 11.903] | < 0.001 | *** |
| AV-cued AV | Visual evidence | 9.237 [8.199, 10.275] | < 0.001 | *** |
| Auditory-cued AV | Previous response | -0.336 [-0.451, -0.220] | < 0.001 | *** |
| Visual-cued AV | Previous response | 0.663 [0.516, 0.810] | < 0.001 | *** |
| AV-cued AV | Previous response | 0.269 [0.130, 0.408] | < 0.001 | *** |
| Auditory-cued AV | Previous auditory direction | -0.021 [-0.146, 0.104] | 0.745 | n.s. |
| Visual-cued AV | Previous auditory direction | -0.069 [-0.209, 0.071] | 0.333 | n.s. |
| AV-cued AV | Previous auditory direction | -0.096 [-0.232, 0.039] | 0.163 | n.s. |
| Auditory-cued AV | Previous visual direction | -0.230 [-0.356, -0.105] | < 0.001 | *** |
| Visual-cued AV | Previous visual direction | -0.402 [-0.565, -0.238] | < 0.001 | *** |
| AV-cued AV | Previous visual direction | -0.189 [-0.341, -0.037] | 0.015 | * |

Several of these trial-level effects diverged from the corresponding PSE-based analyses reported above. The GLMM indicated an attractive effect of previous response in the visual-only and visual-cued AV conditions, in contrast to the repulsive response-history effect indicated by the PSE analysis in the same conditions (section 2.1, Figure 2). The stimulus-direction comparison was more complex than a simple magnitude difference: the direction-based PSE analysis indicated a significant **attractive** effect confined to audition, with no significant effect in vision, whereas the GLMM identified significant **repulsive** stimulus-direction effects in both modalities — the sign in audition is reversed between the two analyses, not just the magnitude. Together, these trial-level results indicate that choice in this task is shaped by dissociable, and in some cases opposing, contributions of response history and stimulus history that differ across modality and cue condition. We consider the source of these discrepancies with the PSE-based analyses, and their implications for the mechanisms underlying serial dependence in this task, in the Discussion.

## 3 Discussion

In this study, we examined how prior trial history influenced response bias in auditory, visual, and audiovisual motion direction discrimination, using both an aggregate psychometric (PSE) approach and a trial-level generalized linear mixed model (GLMM) that isolated the independent contributions of previous stimulus direction and previous response while controlling for current evidence and other trial-level covariates. Both approaches agreed that serial dependence in this task manifested as a shift in response bias, indicating that prior trial history alters decision criteria of the motion judgment. Beyond this point of agreement, the two approaches diverged in informative ways. The PSE-based analysis (sections 2.1–2.2) suggested a relatively clean dissociation, with auditory conditions showing attractive serial dependence and visual conditions showing repulsive serial dependence. The GLMM (section 2.3), which estimated the independent contributions of previous response and previous stimulus direction simultaneously rather than marginally, indicated a more granular structure: a repulsive, stimulus-driven history effect that was strongest and most consistent for vision, and an attractive, response-driven history effect that emerged specifically in conditions where visual information was task-relevant.

This process-level dissociation is broadly consistent with previous work showing that serial dependence is not a unitary phenomenon, reflecting attractive averaging of redundant sensory information under some conditions and repulsive, adaptation-like effects under others, particularly for dynamic stimuli such as motion (Fritsche et al. [2017], Alais et al. [2017]). Recent evidence further suggests that these effects may be supported by partially dissociable modality-specific mechanisms, with auditory and visual serial dependence differing in their feature and spatial selectivity, which provides evidence that history effects do not need to arise from a single modality-general process ([Togoli et al., 2026]). Conversely, other work suggests that serial dependence can also transfer across sensory modalities and be modulated by cross-modal attention, with EEG evidence suggesting that both unimodal and cross-modal effects emerge during perceptual processing ([Fornaciai et al., 2026]). Because motion perception depends on sensitivity to change over time, a purely attractive strategy that averages successive estimates would be computationally counterproductive; a repulsive, decorrelating strategy is more consistent with established motion aftereffects (MAEs), in which recent exposure to motion in one direction biases subsequent perception in the opposite direction ([Glasser et al., 2011, Kanai and Verstraten, 2005, Kohn, 2007, Huk et al., 2001]). Our GLMM results indicate that this repulsive, adaptation-like signature is carried specifically by the stimulus-direction history term, and is most robust for vision: previous visual direction produced a significant repulsive effect not only in the visual-only condition, but across every cued audiovisual condition, regardless of which modality participants were instructed to attend. Previous auditory direction, by contrast, produced a repulsive effect only in the auditory-only condition and was not a reliable predictor in any cued audiovisual condition. This asymmetry suggests that the repulsive, stimulus-driven component of serial dependence in this task is disproportionately visual in origin, persisting even when audition is the behaviorally relevant modality, and is normally masked once visual information is simultaneously present.

This visual specificity fits naturally within the Modality Appropriateness Hypothesis [Welch and Warren, 1980] and our prior demonstration of visual dominance in spatial direction discrimination [Tiesman et al., 2026]. The same asymmetry was evident in our current-evidence weights, which were consistently steeper for visual than auditory motion in the unisensory experiment (Visual*−* Auditory contrast: *β* = 0.76, *p <* .001) and across every cued audiovisual condition, where visual evidence was at least as strong a predictor of choice as auditory evidence even when audition was the cued modality (Figure 4; Tables 2 and 4). A visual system specialized for spatial, fine-grained discrimination is well-positioned to support a fast, low-level adaptation mechanism that discounts recently seen motion directions in the service of change detection [Clifford, 2002, Levinson and Sekuler, 1976]. The comparatively weak and context-dependent auditory stimulus-history effect is consistent with audition’s lesser role in fine spatial discrimination, and, by extension, in a spatially-referenced adaptation process for motion direction specifically [Kohn, 2007, Glasser et al., 2011, Huk et al., 2001].

**Table 3.** Unisensory GLMM: pairwise (Visual*−* Auditory) comparisons of current-evidence weight and trial-history effects. Estimates are GLMM log-odds coefficients (95% CI). N=57 participants, 19404 trials. *p*-values Holm-Bonferroni corrected within this table (correlated pairwise contrasts from one fitted model).

| Condition A | Condition B | Term | Estimate [95% CI] | $p$ | Sig. |
| --- | --- | --- | --- | --- | --- |
| Visual only | Auditory only | Current evidence | 0.760 [0.390, 1.131] | < 0.001 | *** |
| Visual only | Auditory only | Previous response | 0.772 [0.630, 0.914] | < 0.001 | *** |
| Visual only | Auditory only | Previous stimulus direction | 0.024 [-0.128, 0.175] | 0.761 | n.s. |

**Table 4.** Cued-AV GLMM: pairwise cue-condition comparisons of current-evidence weight and trial-history effects. Estimates are GLMM log-odds coefficients (95% CI). N=33 participants, 22736 trials. *p*-values Holm-Bonferroni corrected within this table (correlated pairwise contrasts from one fitted model).

| Condition A | Condition B | Term | Estimate [95% CI] | $p$ | Sig. |
| --- | --- | --- | --- | --- | --- |
| Visual-cued AV | Auditory-cued AV | Auditory evidence | -3.792 [-4.385, -3.199] | < 0.001 | *** |
| AV-cued AV | Auditory-cued AV | Auditory evidence | -1.561 [-2.085, -1.038] | < 0.001 | *** |
| Visual-cued AV | AV-cued AV | Auditory evidence | -2.231 [-2.875, -1.587] | < 0.001 | *** |
| Visual-cued AV | Auditory-cued AV | Visual evidence | 6.819 [6.220, 7.418] | < 0.001 | *** |
| AV-cued AV | Auditory-cued AV | Visual evidence | 5.229 [4.704, 5.754] | < 0.001 | *** |
| Visual-cued AV | AV-cued AV | Visual evidence | 1.590 [0.940, 2.240] | < 0.001 | *** |
| Visual-cued AV | Auditory-cued AV | Previous response | 0.999 [0.812, 1.185] | < 0.001 | *** |
| AV-cued AV | Auditory-cued AV | Previous response | 0.604 [0.424, 0.785] | < 0.001 | *** |
| Visual-cued AV | AV-cued AV | Previous response | 0.394 [0.192, 0.597] | < 0.001 | *** |
| Visual-cued AV | Auditory-cued AV | Previous auditory direction | -0.048 [-0.236, 0.139] | 1.000 | n.s. |
| AV-cued AV | Auditory-cued AV | Previous auditory direction | -0.076 [-0.260, 0.109] | 1.000 | n.s. |
| Visual-cued AV | AV-cued AV | Previous auditory direction | 0.027 [-0.167, 0.222] | 1.000 | n.s. |
| Visual-cued AV | Auditory-cued AV | Previous visual direction | -0.171 [-0.377, 0.035] | 0.515 | n.s. |
| AV-cued AV | Auditory-cued AV | Previous visual direction | 0.042 [-0.156, 0.239] | 1.000 | n.s. |
| Visual-cued AV | AV-cued AV | Previous visual direction | -0.213 [-0.436, 0.010] | 0.368 | n.s. |

**Figure 4.**
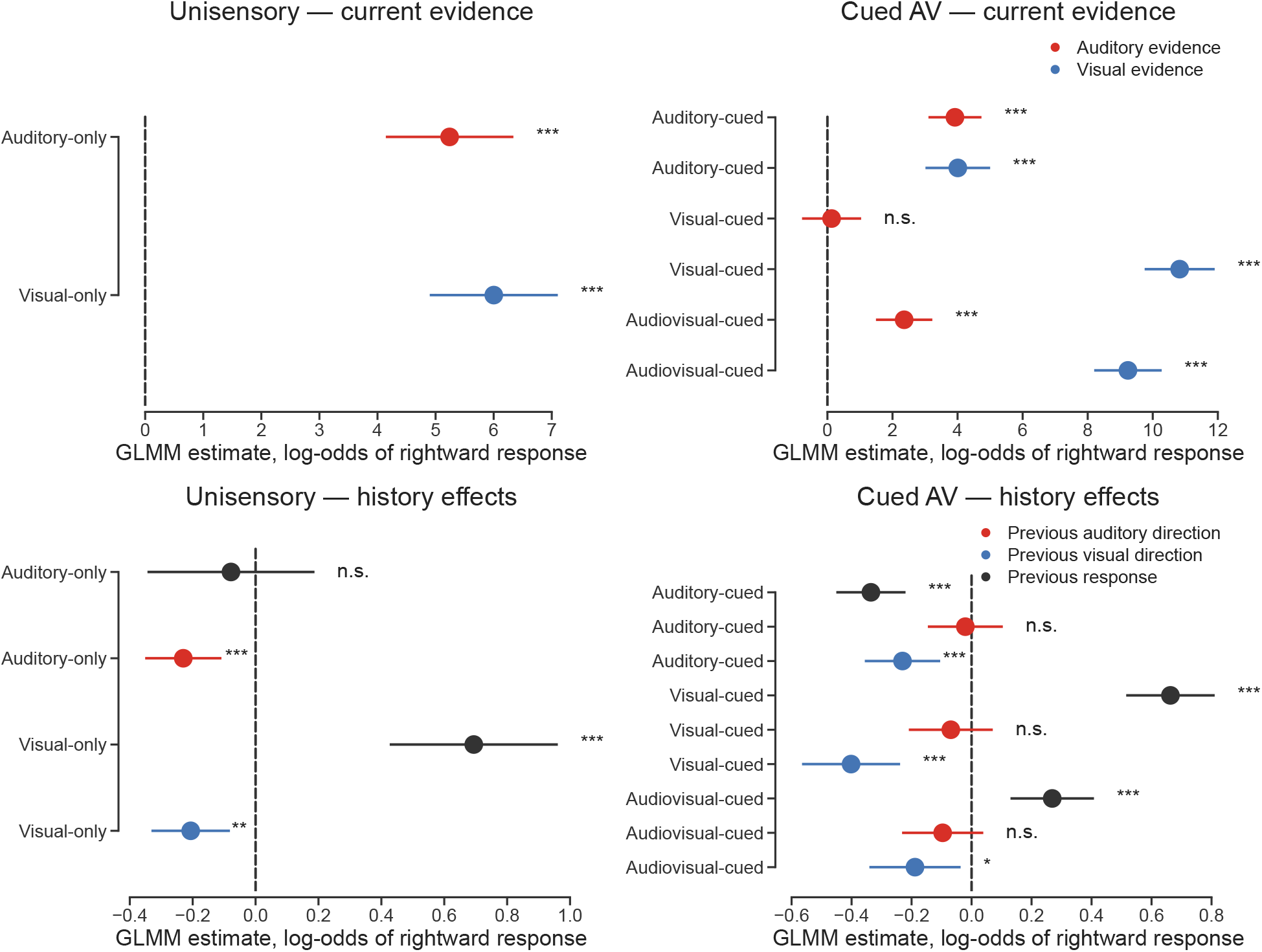
Trial-level GLMM estimates of current-evidence and trial-history effects on motion-direction judgments. All estimates are fixed-effect coefficients in log-odds units of a rightward response, with horizontal bars showing 95% confidence intervals; the dashed vertical line marks zero. For current-evidence terms, larger positive values indicate stronger stimulus control over choice. For history terms, positive values indicate an attractive effect and negative values a repulsive effect. **Top left:** Current-evidence weights by unisensory condition. **Top right:** Auditory- and visual-evidence weights by cue condition in the cued audiovisual experiment. **Bottom left:** Previous-response and previous-stimulus-direction effects by unisensory condition. **Bottom right:** Previous-response, previous-auditory-direction, and previous-visual-direction effects by cue condition. Significance markers refer to tests of each coefficient against zero: * *p <* 0.05, ** *p <* 0.01, *** *p <* 0.001, n.s. *p >* 0.05.

The response-history (choice repetition) component told a different story, and is where the PSE and GLMM results are most directly in tension. The GLMM indicated an attractive effect of the previous response in the visual-only, visual-cued, and audiovisual-cued conditions, but a repulsive effect in the auditory-cued condition and no reliable effect in the auditory-only condition – a pattern that tracks visual task-relevance more than a fixed modality. This is the opposite sign from the visual-only and visual-cued PSE results, which indicated repulsive response-history effects, and the opposite pattern from the auditory PSE results, which indicated attractive response-history effects. We believe this divergence arises because the PSE approach estimates the marginal influence of previous response by fitting separate psychometric curves to trials grouped by that single factor, without simultaneously conditioning on previous stimulus direction or on trial-level nuisance factors (current and previous coherence, catch trials, trial order) that the GLMM includes explicitly. Because previous response and previous stimulus direction are correlated by construction – participants are, on average, more often correct than not – a marginal split on one factor necessarily carries some of the influence of the other. The PSE approach also summarizes each participant with an independently fit psychometric curve before averaging across participants, a two-stage procedure that can behave differently from the GLMM’s single-stage, trial-weighted pooling. We therefore treat the GLMM’s simultaneous, trial-level estimates as the more mechanistically interpretable account of response-history and stimulus-history effects individually, and treat the PSE results as an aggregate, and in several conditions confounded, blend of the two.

Read this way, the overall pattern suggests two dissociable and only partially modality-specific mechanisms rather than a single auditory-versus-visual axis ([Fritsche et al., 2017, Togoli et al., 2026]). Previous stimulus direction, particularly visual stimulus direction, exerted a consistent repulsive influence, most parsimoniously interpreted as a sensory-level, adaptation-like process that shifts perceived motion away from the recently seen direction independent of what the participant last decided ([Kohn, 2007, Glasser et al., 2011, Huk et al., 2001]). Previous response, by contrast, exerted an attractive influence gated by visual task-relevance rather than tied to a single modality – a pattern more consistent with a decisional or post-perceptual locus (e.g., criterion drift or response inertia) engaged specifically when the visual system is driving the choice ([Fritsche et al., 2017]). These two processes are not mutually exclusive, and our results suggest they operate concurrently and, in some conditions, in direct opposition.

This opposition offers a clearer account of the null PSE shift observed in the audiovisual-cued condition (section 2.1, Figure 2) than a simple cancellation via auditory attractive and visual repulsive effects. Rather than reflecting an absence of serial dependence, the GLMM indicated that the audiovisual-cued condition contained two simultaneously significant, oppositely-signed effects – an attractive response-history effect and a repulsive previous-visual-direction effect (section 2.3, Figure 4) – whose sum approximates zero at the level of an aggregate PSE shift. The PSE result correctly detects no net bias, while the GLMM reveals that this apparent absence is the numerical result of two real, opposing history-dependent processes rather than the absence of history-dependence itself.

A plausible neural account can be envisioned for these findings. It can be posited that the repulsive, stimulus-driven component reflects adaptation-like tuning changes within motion-selective visual populations (e.g., hMT+) [Kohn and Movshon, 2004], largely insulated from task instruction, while the attractive, response-driven component reflects a higher-level decisional signal, likely arising in frontoparietal cortices (e.g., dorsolateral prefrontal cortex, posterior parietal cortex) ([Kohn, 2007, Huk et al., 2001, Glasser et al., 2011]). This network is recruited when the visual channel is guiding behavior, potentially propagating via feedback to shape choice on the subsequent trial ([Gold and Shadlen, 2007, Heekeren et al., 2008]). Under this framework, multisensory integration in this task involves not only the fusion of concurrent auditory and visual evidence, but the concurrent, and sometimes competing, influence of a stimulus-level history process and a decision-level history process, consistent with broader accounts of multisensory perception as integrating both current evidence and history-dependent priors ([Shams and Beierholm, 2022, Rohe and Noppeney, 2015, Schwiedrzik et al., 2014]).

Several limitations qualify these conclusions. Most directly, our PSE- and GLMM-based estimates diverge for several effects, so some of our conclusions depend on which analytic approach is taken to more accurately isolate a given history component. We have argued for the trial-level estimates on the grounds that they condition on both history factors simultaneously, but we did not formally verify that this accounts for the divergence, and doing so is an important direction for follow-up work. We nonetheless regard the trial-level pattern as defensible: it was consistent across two independent experiments and across cue conditions within each, which argues against its arising from any single model fit. The comparison is further complicated in the cued audiovisual experiment by a difference in sample, as the PSE analyses treated each condition independently and so retained participants with partial data, whereas the GLMM required valid trials in all three cue conditions.

Relatedly, our GLMM coefficients describe the average influence of trial history across pooled trials, and the scatter of individual ΔPSE values in Figures 2 and 3 makes clear that participants varied considerably in both the magnitude and the direction of their history effects. Although participant-level random effects accommodate this variability during estimation, we did not characterize it directly, and the group-level coefficients we report should not be taken to imply that every participant expressed the same pattern; whether this variability reflects stable individual differences in history-weighting or trial-to-trial noise is an open question that a dedicated, individual-differences design would be better suited to address.

Other limitations concern the scope of what we tested. Our analysis was restricted to n-1 trials, leaving longer-range dependencies unexplored, and — particularly in the audiovisual conditions — we cannot rule out that participants’ compliance with the attentional cue varied. Additionally, while stimulus statistics were tightly controlled, it remains unclear whether these effects would generalize to more naturalistic conditions or to clinical or developmental populations, where multisensory integration and prior-weighting are known to differ [Noel et al., 2018, Wallace et al., 2020]. Finally, complementary neurophysiological measures such as EEG could help directly dissociate early sensory-adaptation signatures from later criterion- or decision-related shifts, rather than inferring this distinction indirectly from behavior.

In summary, our results indicate that serial dependence in auditory, visual, and audiovisual motion judgments is not well characterized as a single attractive-versus-repulsive, auditory-versus-visual dissociation. Instead, trial-level modeling reveals at least two separable processes: a repulsive, largely visual, stimulus-driven history effect that persists across attentional contexts, and an attractive, response-driven history effect selectively engaged when vision is task-relevant. The net bias captured by conventional PSE-based measures – including the apparent absence of serial dependence in the audiovisual-cued condition – reflects the aggregate balance of these underlying components rather than their individual presence or absence. Future work should formally reconcile aggregate and trial-level estimates of serial dependence, and use neurophysiological measures to test whether these statistically dissociable components correspond to distinct sensory- and decision-level neural mechanisms.

## 4 Methods

### 4.1 Participants

Sixty-three participants (median age = 24, 42 females) took part in the first experiment and sixty participants (median age = 19, 39 females) took part in the second experiment. Participants consisted of Vanderbilt undergraduate students and adults from the Nashville metropolitan area with normal or corrected-to-normal vision, normal hearing, and no known neurological disorders. Undergraduate participants were compensated with course credits, and non-undergraduate participants were compensated monetarily at a rate of $20/hour. Six participants in the first experiment and twelve participants in the second experiment were excluded from behavioral analyses because they did not complete all task conditions, leaving a final analyzed sample of 57 participants for the first experiment and 48 participants for the second experiment. Each participant gave their informed consent before being allowed to participate, and all recruitment procedures were in accordance with the Vanderbilt University Psychology Department Guidelines. Basic participant demographics including sex and age were also recorded during consent. All recruitment and experimental procedures were approved by the Vanderbilt University Institutional Review Board and were carried out in accordance with the Declaration of Helsinki.

### 4.2 Stimuli

In the task, participants were presented with visual, auditory, or audiovisual stimuli. All stimuli were generated in MATLAB R2022b (The MathWorks, Inc., Natick, MA) and presented using PsychToolbox version 3 ([Brainard, 1997, Kleiner et al., 2007]). Both unisensory (auditory only and visual only) and multisensory (auditory and visual presented at the same time) stimuli were included in the experiment. Stimuli had a duration of 700 milliseconds, and were independently generated for each trial, and therefore not “frozen.”

Visual stimuli reflected a Movshon/Newsome-type motion algorithm and were displayed on a 1920×1080 cathode ray tube monitor positioned at eye-level approximately 120 cm from the participant ([Britten et al., 1992, Newsome and Paré, 1988]). The stimuli were random-dot kinematograms (RDKs) with 150, 3 pixel white dots (subtending <0.3° of visual space) placed randomly in a square aperture subtending a width and height of 14° visual space ([Tiesman et al., 2026]). The aperture size reflected the simulated displacement of the auditory stimulus at the same speed of 20°/s and a stimulus duration of 700 milliseconds. The proportion of dots that moved in a signal direction, either left or right, corresponded to the coherence value for the visual stimulus, while the remaining dots moved in random directions. Larger coherence values indicated a greater strength of motion, due to more dots within the aperture moving in the same signal direction.

Auditory stimuli consisted of 3 milliseconds of silence followed by a 700 millisecond broad-band white noise signal (10ms rise and fall) sampled at a rate of 44100 samples per second. The signal moved at a speed of 20°/s, was embedded in partially correlated noise and played through two speakers mounted on either side of the monitor (54 cm apart) centered horizontally with the visual stimulus, on the azimuth for the participant. The auditory stimulus consisted of four different components. Independent white noise streams were presented through both speakers (100% amplitude; inter-signal correlation = 0; N1 and N2). On top of this, a combined white noise signal was presented through both speakers (100% amplitude, inter-signal correlation = 1; N3). The fourth signal stream contained the apparent motion cue, in which the sound’s amplitude faded between the two speakers from 100% to 0% over the course of one trial (N4) to create binaural motion cues. Coherence values for the auditory stimulus were the signal level, or N4 component, to noise level, N1-N3 components, ratio. Similar to the visual stimulus, larger coherence values indicated a greater strength of motion, as the N4 motion signal was louder compared to the noise. The two speakers playing the motion stimulus generated 66 dB SPL measured at the ear-level of the participant ([Tiesman et al., 2026]).

A Minolta Chroma Meter CS-100 and a sound level meter (PCB Piezotronics, Model 378C01) were used to verify the luminance and sound intensity levels, respectively. The durations of all visual and auditory stimuli were confirmed using a Hameg 507 oscilloscope (Hameg Instruments, Mainhausen, Germany) with a photovoltaic cell and microphone. All stimulus parameters of this paper were adapted from [Tiesman et al., 2026].

### 4.3 Paradigm

Participants were presented with a series of two-alternative forced choice motion discrimination tasks in either unisensory or audiovisual conditions. All procedures of this paper were adapted from [Tiesman et al., 2026].

#### Experiment 1

In this experiment, participants were presented with two unisensory conditions consisting of either the auditory stimulus (auditory-only condition) or the visual stimulus (visual-only condition). The order of the unisensory conditions was randomized for each participant. Prior to starting, participants were instructed to maintain fixation on a central point and to report the general direction of a motion stimulus (right or left) via a response box as quickly and accurately as possible. Upon starting, all trial directions were random and had an equal probability of being left or right. Assignment of trial coherence was also random, with each unisensory condition containing 40 trials per coherence value, along with 20 noise trials, for a total of 180 trials. No feedback on performance was given. A new trial was only initiated after the stimulus was presented and the additional response period had elapsed, regardless of whether the participant had responded.

On each trial of the experiment, participants were able to respond while the stimulus was being presented or during a subsequent response period. The stimulus was presented for 700 milliseconds, and the time of the subsequent response period was randomly selected from a uniform distribution of 1.5 and 2.5 seconds. The fixation cross in the middle of the CRT screen remained visible for the duration of the trial. A new trial was only initiated after the stimulus was presented and the additional response period had elapsed and was initiated whether or not the participant responded.

#### Experiment 2

Participants were initially presented with two adaptive unisensory staircase conditions (auditory-only and visual-only) for the purpose of finding coherence thresholds and sensory weights. Then, in the main part of the experiment, participants were presented with three audiovisual conditions. Each audiovisual condition had the same number of trials and stimuli, although the task instructions given prior to the start of the condition differed. Participants were instructed to report the direction of motion of only the auditory stimulus (auditory-cued condition), only the visual stimulus (visual-cued condition), or both the auditory and visual stimuli (audiovisual-cued condition). All other instructions were the same as those in experiment 1. Presentation order of the auditory cued and visual cued conditions was randomized for each participant. In each condition, each coherence level had 48 trials, with an additional 20 complete auditory and visual noise trials interspersed throughout, for a total of 212 trials per condition. Each set of 48 trials was evenly split into leftward and rightward trials. Additionally, the sets consisted of 40 congruent trials and 8 incongruent trials, evenly split between leftward and rightward trials. The auditory and visual stimuli had the same direction in congruent trials and the opposite directions in the incongruent trials, and participants were not told about the congruency of the motion. Based on a pilot study, the greater proportion of incongruent trials significantly increased the difficulty of the task. The current proportion of congruent to incongruent trials was to ease difficulty while also interrogating the incongruent trial effects. Experiment 2’s trial length and structure was identical to that of Experiment 1.

### 4.4 Data Analysis

Responses were collected across a range of motion coherence to assess participant performance, and these were fit to a cumulative Gaussian distribution. The x-value for a 0.5 proportion rightward response was used as the point of subjective equality (PSE), a measure used to examine directional bias across participants for our task. For this measure, a negative PSE indicated a rightward bias, whereas a positive PSE indicated a leftward bias.

Outliers in point of subjective equality values were assessed separately for each condition using both the median absolute deviation (MAD)–based robust z-score (> |3.5|) and the Tukey interquartile range (IQR) method (±1.5×IQR). PSE values flagged by either criterion were removed from subsequent group-level analyses. Importantly, only the affected condition(s) were excluded for a given participant; remaining conditions for that participant were retained to maximize valid data contributions. Subsequent paired within-group comparisons of PSE were performed using Wilcoxon signed-rank tests. Holm-Bonferroni correction was applied to control for multiple comparisons where appropriate.

### 4.5 Generalized linear mixed-effects modeling

Trial-level choices were modeled using generalized linear mixed-effects models (GLMMs) with a binomial distribution and logit link, fit by maximum likelihood using the Laplace approximation (MATLAB R2022b, Statistics and Machine Learning Toolbox, fitglme). The outcome variable was the binary response on each trial (1 = rightward, 0 = leftward). Fixed effects included signed current sensory evidence (coherence-weighted, signed for direction), a nuisance term for unsigned coherence, trial number (z-scored), and trial-history predictors: the participant’s response on the previous trial, the stimulus direction on the previous trial, the coherence of the previous trial, and whether the previous trial was a catch (zero-coherence) trial. History predictors were effect-coded (±0.5) rather than dummy-coded so that main-effect coefficients reflected the full leftward-to-rightward contrast independent of interaction terms.

For the unisensory model, trials were coded by modality (auditory-only vs. visual-only, auditory-only as the reference level), and all evidence and history terms were allowed to interact with modality to yield modality-specific coefficients. The cued audiovisual model followed the same logic, with auditory-cued, visual-cued, and audiovisual-cued trials (auditory-cued as reference) and separate auditory- and visual-evidence terms, each interacting with cue condition. Because the cued audiovisual model estimates cue-condition-specific coefficients within a single fitted model, only participants with valid trials in all three cue conditions were included; this criterion yielded 33 of the 48 behaviorally-analyzed participants for the cued audiovisual GLMM. Random effects included participant-specific intercepts and slopes for current evidence and (where applicable) response- and stimulus-history terms, capturing individual differences in overall bias, evidence sensitivity, and history weighting; the random-effects structure was simplified from a maximal specification only as needed to reach convergence, with the same structure used across nested models to preserve valid likelihood-ratio comparisons. The contribution of trial-history terms was assessed by comparing a full model against a reduced model containing only current-evidence terms via a likelihood-ratio test, and condition-specific effects and pairwise condition contrasts were computed as linear combinations of fixed-effect coefficients, with standard errors derived from the full coefficient covariance matrix. Coefficients are reported in log-odds units, with positive values indicating a rightward bias and negative values a leftward bias; for history terms, positive coefficients indicate an attractive (repetition-promoting) effect and negative coefficients a repulsive (alternation-promoting) effect.

## 5 Author Contributions

A.J.T. contributed to study conception, study design, data collection, data analysis, figure generation, manuscript writing, and manuscript editing. K.S. contributed to study design, data collection, data analysis, figure generation, manuscript writing, and manuscript editing. H.B. contributed to data collection, data analysis, and manuscript editing. R.R. and

M.T.W. contributed to study conception, study design, study supervision, and manuscript editing. All authors reviewed the manuscript.

## 7 Acknowledgments

The authors would like to thank Elizabeth Brandon and Mckenzie King for their assistance with participant recruitment.

## References

D. Alais and D. Burr. The ventriloquist effect results from near-optimal bimodal integration. Curr Biol, 14(3):257–62, 2004.

David Alais, Johahn Leung, and Erik Van der Burg. Linear Summation of Repulsive and Attractive Serial Dependencies: Orientation and Motion Dependencies Sum in Motion Perception. Journal of Neuroscience, 37(16):4381–4390, April 2017. ISSN 0270-6474, 1529-2401. doi: 10.1523/JNEUROSCI.4601-15.2017. URL. https://www.jneurosci.org/content/37/16/4381.

P. W. Battaglia, R. A. Jacobs, and R. N. Aslin. Bayesian integration of visual and auditory signals for spatial localization. J Opt Soc Am A Opt Image Sci Vis, 20(7):1391–7, 2003.

David H. Brainard. The Psychophysics Toolbox. Spatial Vision, 10(4):433–436, 1997.

K. H. Britten, M. N. Shadlen, W. T. Newsome, and J. A. Movshon. The analysis of visual motion: a comparison of neuronal and psychophysical performance. J Neurosci, 12(12):4745–65, 1992.

Guido Marco Cicchini, Giovanni Anobile, and David C. Burr. Compressive mapping of number to space reflects dynamic encoding mechanisms, not static logarithmic transform. Proceedings of the National Academy of Sciences, 111(21):7867–7872, May 2014. doi: 10.1073/pnas.1402785111. URL https://www.pnas.org/doi/full/10.1073/pnas.1402785111.

Guido Marco Cicchini, Kyriaki Mikellidou, and David Charles Burr. Serial Dependence in Perception. Annual Review of Psychology, 75:129–154, 2024. ISSN 1545-2085. doi: 10.1146/annurev-psych-021523-104939. URL https://www.annualreviews.org/content/journals/10.1146/annurev-psych-021523-104939.

Colin W. G. Clifford. Perceptual adaptation: motion parallels orientation. Trends in Cognitive Sciences, 6(3):136–143, March 2002. ISSN 1364-6613. doi: 10.1016/S1364-6613(00)01856-8. URL https://www.sciencedirect.com/science/article/pii/S1364661300018568.

M. O. Ernst and M. S. Banks. Humans integrate visual and haptic information in a statistically optimal fashion. Nature, 415(6870):429–33, 2002.

Jason Fischer and David Whitney. Serial dependence in visual perception. Nature Neuroscience, 17(5):738–743, May 2014. ISSN 1546-1726. doi: 10.1038/nn.3689. URL https://www.nature.com/articles/nn.3689.

Michele Fornaciai, Irene Togoli, Samuel Binisti, and Olivier Collignon. Serial dependence generalizes across the senses, March 2026. URL https://www.biorxiv.org/content/10.64898/2026.03.16.712008v1. xISSN: 2692-8205 Pages: 2026.03.16.712008 Section: New Results.

T. C. Freeman, J. Leung, E. Wufong, E. Orchard-Mills, S. Carlile, and D. Alais. Discrimination contours for moving sounds reveal duration and distance cues dominate auditory speed perception. PLoS One, 9(7):e102864, 2014.

Matthias Fritsche, Pim Mostert, and Floris P. de Lange. Opposite Effects of Recent History on Perception and Decision. Current biology: CB, 27(4):590–595, February 2017. ISSN 1879-0445. doi: 10.1016/j.cub.2017.01.006.

Davis M. Glasser, James M. G. Tsui, Christopher C. Pack, and Duje Tadin. Perceptual and neural consequences of rapid motion adaptation. Proceedings of the National Academy of Sciences, 108(45):E1080–E1088, November 2011. doi: 10.1073/pnas.1101141108. URL https://www.pnas.org/doi/10.1073/pnas.1101141108.

Joshua I. Gold and Michael N. Shadlen. The neural basis of decision making. Annual Review of Neuroscience, 30: 535–574, 2007. ISSN 0147-006X. doi: 10.1146/annurev.neuro.29.051605.113038.

Shinya Harada, Ryo Teraoka, Naoki Kuroda, and Wataru Teramoto. Sound-induced visual motion perception in older adults: aging enhances audiovisual motion integration. Experimental Brain Research, 244(5):86, April 2026. ISSN 1432-1106. doi: 10.1007/s00221-026-07286-x. URL https://doi.org/10.1007/s00221-026-07286-x.

Takuma Hashimoto and Shogo Makioka. Serial dependence in numerosity perception generalizes across different sensory modalities: evidence from sequential numerosity comparison. Acta Psychologica, 265:106691, May 2026. ISSN 0001-6918. doi: 10.1016/j.actpsy.2026.106691. URL https://www.sciencedirect.com/science/article/pii/S0001691826004920.

Hauke R. Heekeren, Sean Marrett, and Leslie G. Ungerleider. The neural systems that mediate human perceptual decision making. Nature Reviews. Neuroscience, 9(6):467–479, June 2008. ISSN 1471-0048. doi: 10.1038/nrn2374.

A. C. Huk, D. Ress, and D. J. Heeger. Neuronal basis of the motion aftereffect reconsidered. Neuron, 32(1):161–172, October 2001. ISSN 0896-6273. doi: 10.1016/s0896-6273(01)00452-4.

Ryota Kanai and Frans A. J. Verstraten. Perceptual manifestations of fast neural plasticity: Motion priming, rapid motion aftereffect and perceptual sensitization. Vision Research, 45(25):3109–3116, November 2005. ISSN 0042-6989. doi: 10.1016/j.visres.2005.05.014. URL https://www.sciencedirect.com/science/article/pii/S0042698905002634.

Christoph Kayser and Herbert Heuer. Multisensory perception depends on the reliability of the type of judgment. Journal of Neurophysiology, 131(4):723–737, April 2024. ISSN 1522-1598. doi: 10.1152/jn.00451.2023.

Anastasia Kiyonaga, Jason M. Scimeca, Daniel P. Bliss, and David Whitney. Serial Dependence across Perception, Attention, and Memory. Trends in Cognitive Sciences, 21(7):493–497, July 2017. ISSN 1879-307X. doi: 10.1016/j.tics.2017.04.011.

M. Kleiner, D. Brainard, and D. Pelli. What’s new in Psychtoolbox-3? page 14, 2007. doi: 10.1177/03010066070360S101. URL https://journals.sagepub.com/doi/pdf/10.1177/03010066070360S101.

Adam Kohn. Visual adaptation: physiology, mechanisms, and functional benefits. Journal of Neurophysiology, 97(5): 3155–3164, May 2007. ISSN 0022-3077. doi: 10.1152/jn.00086.2007.

Adam Kohn and J. Anthony Movshon. Adaptation changes the direction tuning of macaque MT neurons. Nature Neuroscience, 7(7):764–772, July 2004. ISSN 1546-1726. doi: 10.1038/nn1267. URL https://www.nature.com/articles/nn1267.

Eugene Levinson and Robert Sekuler. Adaptation alters perceived direction of motion. Vision Research, 16(7):779–IN7, January 1976. ISSN 0042-6989. doi: 10.1016/0042-6989(76)90189-9. URL https://www.sciencedirect.com/science/article/pii/0042698976901899.

Michael L. Morgan, Gregory C. DeAngelis, and Dora E. Angelaki. Multisensory Integration in Macaque Visual Cortex Depends on Cue Reliability. Neuron, 59(4):662–673, 2008.

Micah M. Murray and Mark T. Wallace, editors. The Neural Bases of Multisensory Processes. CRC Press, Boca Raton, August 2011. ISBN 978-0-429-13010-6. doi: 10.1201/9781439812174.

Michael F. Neelon and Rick L. Jenison. The temporal growth and decay of the auditory motion aftereffect. The Journal of the Acoustical Society of America, 115(6):3112–3123, June 2004. ISSN 0001-4966. doi: 10.1121/1.1687834. URL https://doi.org/10.1121/1.1687834.

W. T. Newsome and E. B. Paré. A selective impairment of motion perception following lesions of the middle temporal visual area (MT). The Journal of Neuroscience: The Official Journal of the Society for Neuroscience, 8(6):2201–2211, June 1988. ISSN 0270-6474. doi: 10.1523/JNEUROSCI.08-06-02201.1988.

Jean-Paul Noel, Ryan A. Stevenson, and Mark T. Wallace. Atypical audiovisual temporal function in autism and schizophrenia: similar phenotype, different cause. European Journal of Neuroscience, 47(10):1230–1241, 2018. ISSN 1460-9568. doi: 10.1111/ejn.13911. URL https://onlinelibrary.wiley.com/doi/abs/10.1111/ejn.13911.

B. Odegaard, D. R. Wozny, and L. Shams. Biases in Visual, Auditory, and Audiovisual Perception of Space. PLoS Comput Biol, 11(12):e1004649, 2015.

Ayberk Ozkirli, Andrey Chetverikov, and David Pascucci. Large-scale mega-analysis indicates that serial dependence deteriorates perceptual decision-making. Nature Human Behaviour, 10(3):568–578, March 2026. ISSN 2397-3374. doi: 10.1038/s41562-025-02362-8. URL https://www.nature.com/articles/s41562-025-02362-8.

David Pascucci, Ömer Dağlar Tanrikulu, Ayberk Ozkirli, Christian Houborg, Gizay Ceylan, Paul Zerr, Mohsen Rafiei, and Árni Kristjánsson. Serial dependence in visual perception: A review. Journal of Vision, 23(1):9, January 2023. ISSN 1534-7362. doi: 10.1167/jov.23.1.9. URL https://pmc.ncbi.nlm.nih.gov/articles/PMC9871508/.

T. Rohe and U. Noppeney. Cortical hierarchies perform Bayesian causal inference in multisensory perception. PLoS Biol, 13(2):e1002073, 2015.

A.M. Schoenhaut, R. Ramachandran, and M.T. Wallace. Contribution of displacement, duration, and velocity on auditory motion direction perception in macaque monkeys. Scientific Reports, 15:28110, 2025.

Caspar M. Schwiedrzik, Christian C. Ruff, Andreea Lazar, Frauke C. Leitner, Wolf Singer, and Lucia Melloni. Untangling Perceptual Memory: Hysteresis and Adaptation Map into Separate Cortical Networks. Cerebral Cortex, 24(5):1152–1164, May 2014. ISSN 1047-3211. doi: 10.1093/cercor/bhs396. URL https://doi.org/10.1093/cercor/bhs396.

Ladan Shams and Ulrik Beierholm. Bayesian causal inference: A unifying neuroscience theory. Neuroscience & Biobehavioral Reviews, 137:104619, June 2022. ISSN 0149-7634. doi: 10.1016/j.neubiorev.2022.104619. URL https://www.sciencedirect.com/science/article/pii/S0149763422001087.

Adam J. Tiesman, Kalina Stoyanova, Ramnarayan Ramachandran, and Mark T. Wallace. Behavioral evidence for visually dominant audiovisual motion integration. Scientific Reports, July 2026. ISSN 2045-2322. doi: 10.1038/s41598-026-62446-x. URL https://www.nature.com/articles/s41598-026-62446-x.

Irene Togoli, Michele Fornaciai, and Domenica Bueti. Different modality-specific mechanisms mediate serial dependence effects in visual and auditory perception. BMC Biology, 24(1):95, March 2026. ISSN 1741-7007. doi: 10.1186/s12915-026-02515-9. URL https://doi.org/10.1186/s12915-026-02515-9.

Erik Van der Burg, Alexander Toet, Anne-Marie Brouwer, and Jan B. F. Van Erp. Serial Dependence of Emotion Within and Between Stimulus Sensory Modalities. Multisensory Research, pages 1–22, September 2021. ISSN 2213-4808. doi: 10.1163/22134808-bja10064.

M. T. Wallace, T. G. Woynaroski, and R. A. Stevenson. Multisensory Integration as a Window into Orderly and Disrupted Cognition and Communication. Annu Rev Psychol, 71:193–219, 2020.

Robert B. Welch and David H. Warren. Immediate perceptual response to intersensory discrepancy. Psychological Bulletin, 88(3):638–667, 1980. ISSN 1939-1455. doi: 10.1037/0033-2909.88.3.638. Place: US.

